# Continuous tracking of gravistimulated roots in a chambered coverslip by confocal microscopy allows first glimpse on mechanoadaptation of cell files during curvature initiation

**DOI:** 10.1101/2021.09.05.459030

**Authors:** Katarzyna Retzer

## Abstract

Mechanical responses of individual cells to plant internal and external stimuli modulate organ movement and ensure plant survival as sessile organism in a constantly changing environment. The root is a complex, three-dimensional object, which continuously modifies its growth path. Autonomous and paratonic root movements are both orchestrated by different signaling pathways, whereby auxin modulated directional growth adaptations, including gravitropic response, were already subject of manifold studies. But we still know very little about how cells adapt upon gravitropic stimulus to initiate curvature establishment, which is required to align root tip growth again along the gravitropic vector. This manuscript shows first insights into cell file movements upon gravitropic stimulus of *Arabidopsis thaliana* roots that initiate curvature establishment. The roots were grown shaded from light and without exogenous sucrose supplementation, both growth conditions that are known to negatively interfere with directed root growth, which allowed a more uniform tracking of root bending by using a confocal microscope with vertical stage.

## Introduction

The root is complex, and not often displayed in manuscript addressing its movements as the three-dimensional object it is, which is composed of different tissues and zones with different roles and mechanical properties. Roots evolved to anchor the plant in the soil and also to absorb water and minerals from it. Furthermore, as all land species, plants developed mechanisms to orient themselves along the gravity vector, whereby roots adapted positive gravitropism which dominantly determines their growth direction ^[1-5]^. Already 1880 Darwin and Darwin suggested in their study ’sThe Power of Movement in Plants’ that gravity perception of roots is localized in the root tip (mechanosensing), a signal is transmitted shootwards (mechanotransmission) shootwards, where it initiates bending (mechanoadaptation) ^[6-8]^. Currently, positive gravitropic response is divided into four steps, a, sensing the gravity vector in the columella, b, biochemical signal transduction, c, differential cell adaptation to induce bending, and d, period of gravitropic signal attenuation to prevent overbending ^[9-11]^.

Roots are very plastic and continuously adapt their appearance and growth direction depending on environmental conditions ^[12-13]^. Directional root growth allows the root to grow towards beneficial areas in the soil and further to avoid toxic environment ^[14]^. Tropistic root growth is described to be driven by asymmetric changes in cell expansion along the root tip to maneuver its path in the most efficient way ^[14]^. Therefore, directional growth of the root results from movement regulation by curvature initiation of the root in the distal elongation zone, also known as transition zone. The dynamics of changes in orientation of the root tip to maintain growth along the gravitropic vector were studied manifold, and it is well described that the root curvature is triggered by the asymmetric auxin distribution between the upper and lower side of root the gravistimulated root ^[2,5,9]^. The efficiency of the response is visible as the angle of the curvature, but how this curvature is established at cellular level is still not fully understood.

## Results and Outlook

### Sample preparation and mounting for microscopy

The root is the belowground organ of the plant, therefore it is, when grown in soil, not easily available for phenotypical studies, and it takes more effort to prepare it for analytical and molecular biological experiments. Therefore, most studies focused on cell biology, analytical studies and microscopy are performed on seedlings grown on agar plates with roots exposed to light, which was manifold shown that it is negatively interfering with cell proliferation, elongation processes, by triggering root escape growth, and even nutrient uptake ^[15-20]^. Furthermore, standard lab conditions include often the usage of media supplemented with sucrose or glucose, which is enhancing root growth, but also interferes with root trait establishment and can further mask phenotypes or cross-talk with stress adaptation ^[15,16, 18-21]^. Sucrose and glucose, especially in combination with direct root illumination result in enhanced deviation of root growth from vertical ^[19-21]^. Therefore, in this study I observed the initiation of gravitropic bending at cellular level of roots, which were grown on 1% agar supplemented medium, but to avoid unnecessary stimuli, exogenous carbon source supplementation was excluded and the so-called D-root device to shade the root from direct illumination was used ^[16,19]^. To track cellular adaptation upon gravitropic stimulus I used a confocal microscope with vertical stage and transferred the seedlings, by cutting out the surrounding growth medium and placing it together in a chambered cover glass ^[22,23]^. *Arabidopsis thaliana Col0* is a well-studied model organism and plenty genetic and molecular tools are established. To be able to clearly track individual cell lines identified to be crucial for the bending process, namely the epidermis and lateral root cap cells (LRC) ^[1]^, I tracked bending of the reporter lines expressing PIN2:GFP^[24]^ and PIN2:mcherry ^[10]^.

### Tracking root bending at cellular level over time shows that LRC keep their position relative to its neighbor epidermis cells upon gravistimulation

After changing the position of the root by 90° by rotating the chamber directly at the vertical stage of the confocal microscope, I immediately focused to the root tip and took pictures over time for 60 minutes and at several positions in the Z-axis of the root, starting at the surface of the root till the center of the root. A 3D modulation of the Z-stacks taken of bending *eir1-4 PIN2::PIN2:mcherry* at timepoint 0’s (Fig1. A) and after 60’ (Fig1.B) of a representative root is shown. It is visible that 60’after gravistimulation the root formed a curvature at its lower side towards the gravitropic vector. Overall, the upper and lower side of the root continued to elongate, and interestingly, the individual LRC cells didn’t change their position relatively to the epidermis cell they were attached to at the beginning of the gravistimulation. The blue arrows in the enlarged segments of the bending root indicate the position of LRC cells at 0’(Fig1. C) and 60’(Fig1. D).

**Figure 1.:**
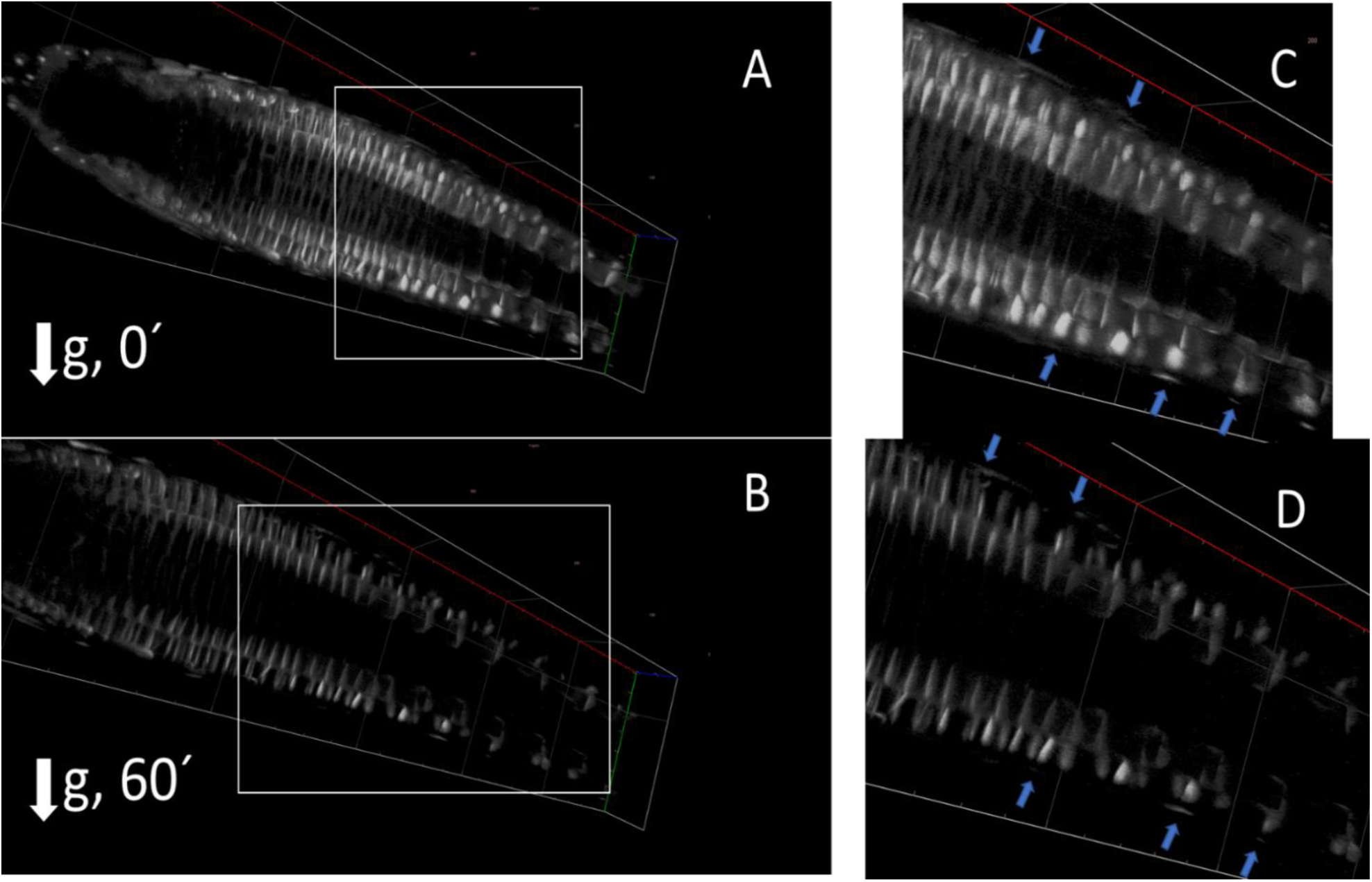
3D alignment of Z-Stacks from the root surface to the root center of a bending root at 0’sand 60’: A, shows 6DAG *eir1-4 PIN2::PIN2:mcherry* at the initiation of the gravistimulus and B, 60’s later. C, and D, show the distal elongation zone enlarged. The blue arrows point towards LRC cells.

The LRC cells seem to adapt morphology by clinging to the epidermis cells, which means for the LRC at the lower side of the bending root to elongate along the curvature in the region of distal elongation zone, which is also visible in Fig2., which shows an optical cut central through a *eir1-4 PIN2::PIN2:mcherry* root, at timepoint 0’s (Fig2. A) and after 60’ (Fig2.B). Furthermore, it looks like the epidermis cells in the distal elongation zone below the elongating LRC are pushed towards the stele, but from this angle of observation and by only comparing the mentioned time points it is not clear how the curvature was initiated.

**Figure 2.:**
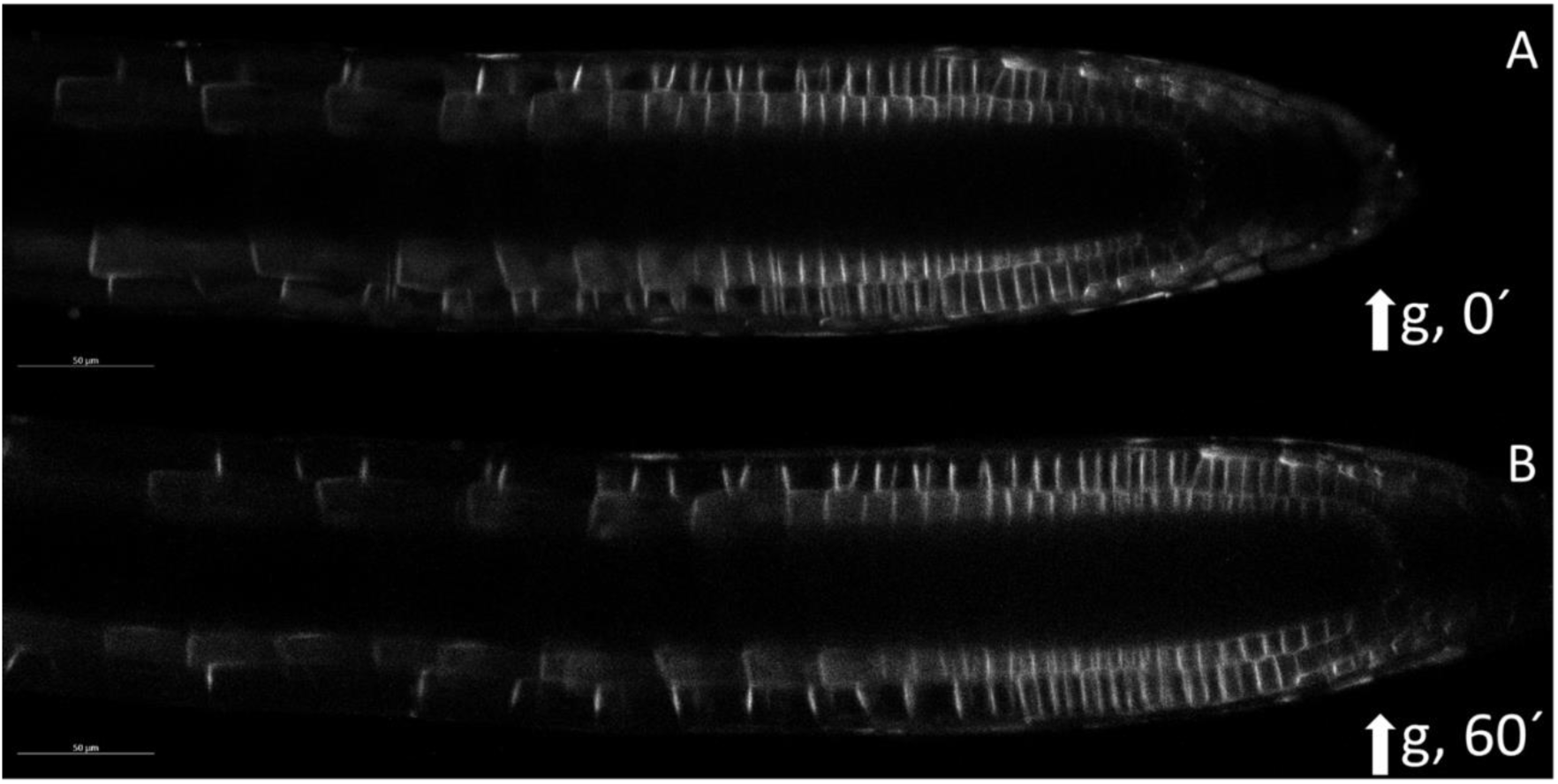
Optical sectioning at the position of the root center of a bending root at 0’sand 60’: A, shows 6DAG *eir1-4 PIN2::PIN2:mcherry* at the initiation of the gravistimulus and B, 60’s later.

### After gravistimulation one epidermal cell file is sliding over its neighbor cell file in opposite direction of the gravitropic vector and thereby initiating the curvature

To capture the events of curvature initiation of gravistimulated roots, I observed continuously for 40’s individual roots at different levels along the Z-axis. A representative root expressing PIN2:GFP to visualize the root cells is shown in Fig. 3. Panels A-D show an optical cut at the position one cell layer above the root center, and panel E-H show the cells central in the root. Movies of the bending root are available under https://youtu.be/EH4ruwqjC4Y (corresponding to panel A-D) and https://youtu.be/38t2C9ovC_s (corresponding to panel E-F). The two optical cuts show different events, which together may explain how the curvature of a graviresponsing root is initiated. Mechnoadaptation of the involved cell files is progressing continuously and requires a three-dimensional observation of the bending root, otherwise the events can be overlooked easily. When tracking the events in the higher positioned cell file, it is already after few minutes visible, as indicated by the orange arrows, that one epidermal cell file is sliding over its neighbor cell file in the opposite direction of the gravitropic vector (Fig. 3. B). The shift in position starts in the meristematic zone and after 20’(Fig. 3. C) till 40’(Fig. 3. D) the movement reaches gradually the distal elongation zone. When observing root bending at the level of the cut through the very center of the root (Fig. 3. E-H), the change in position of this particular cell file relatively to its neighbor cell files is not visible to the same extend as mentioned above. But at this level the change in cell morphology is better visible. Individual cells in the distal elongation zone that are positioned in the middle of the appearing curvature are marked with blue stars. These particular cells appear in the higher positioned optical cut in the region of interest around 20’ after gravistimulation of the root (Fig. 3. C), and are visible in the lower optical segment from the very beginning (Fig. 3. E-H).

**Figure 3.:**
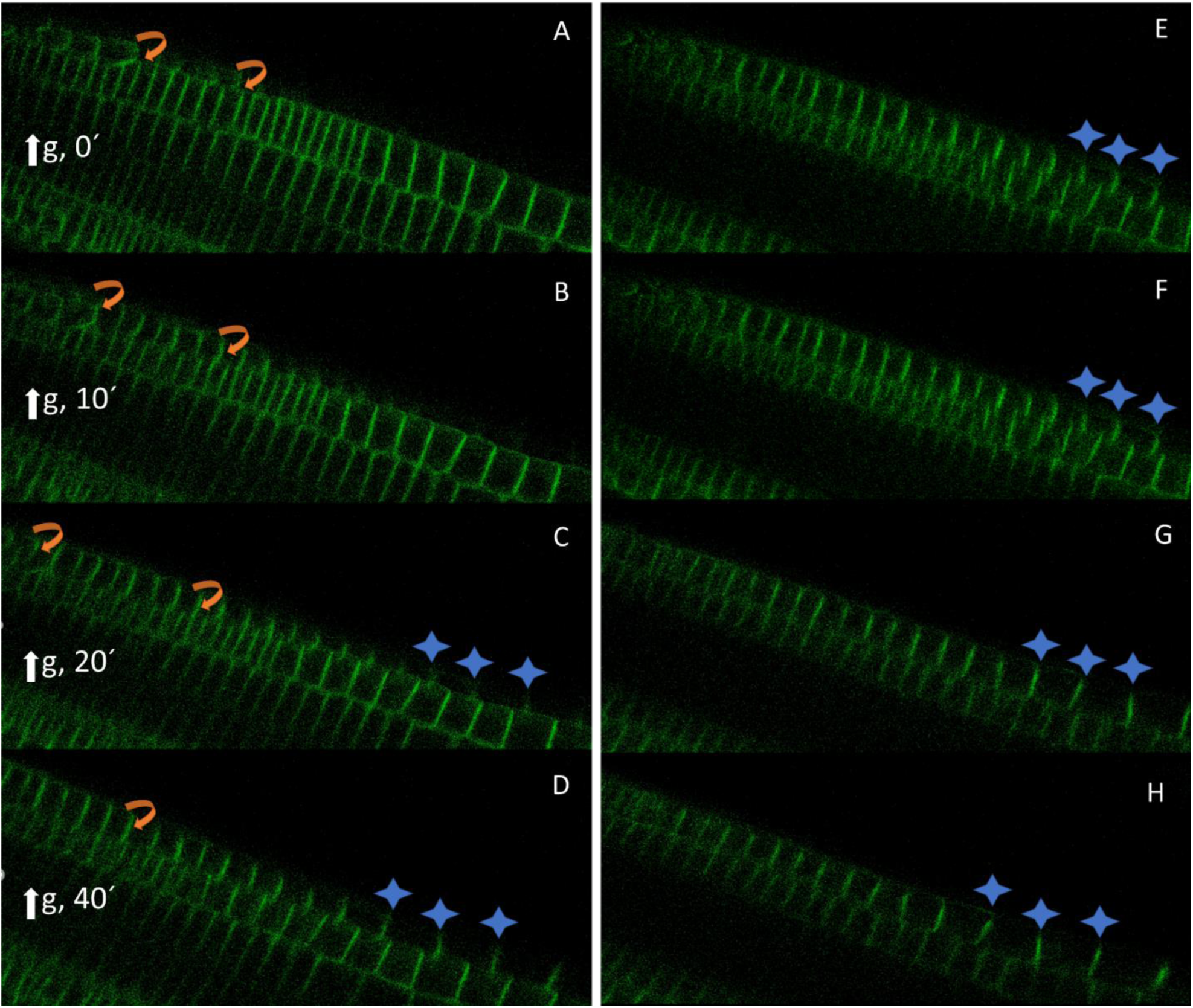
Optical sectioning at two positions along the Z-axis of a bending root over time. 7 DAG *Col0* expressing PIN2:GFP was gravistimulated and continuously observed for 40’s. Panel A-D show response of cell files one cell layer above the center of the root, whereas panel E-H show the section in the center of the root. The orange arrows indicate the rotation of an epidermal cell file over time upon gravistimulation. The blue stars mark particular cells that change their position upon gravistimulation.

By tracking a root of a 4DAG seedling, with root grown in darkness, over 105min after gravistimulation, it is visible that the epidermal cells immediately responding to gravistimulus shift position by cell file rotation starting at the very root tip up to the start of the elongation zone, where they movement induce in the neighbor cell file the adjustment of cell expansion that finally allows root curvature formation (Fig. 4). Selected time points are aligned in Fig 4., and the rotating cell file is marked in red, whereas the neighbor cell file that responds in the elongation zone prior to directional growth adaptation is marked blue. The corresponding movie can be found under https://www.youtube.com/channel/UC9tji6ljLRuo-nQbSun26EA, as well videos of Z-stacks taken at position below the root epidermis surface. In total I obtained continuously with the 20x objective 3 tiles, over 8 Z-stacks every 3μm for 105 min., by averaging every picture twice, which resulted in 49 timepoints.

**Figure 4.:**
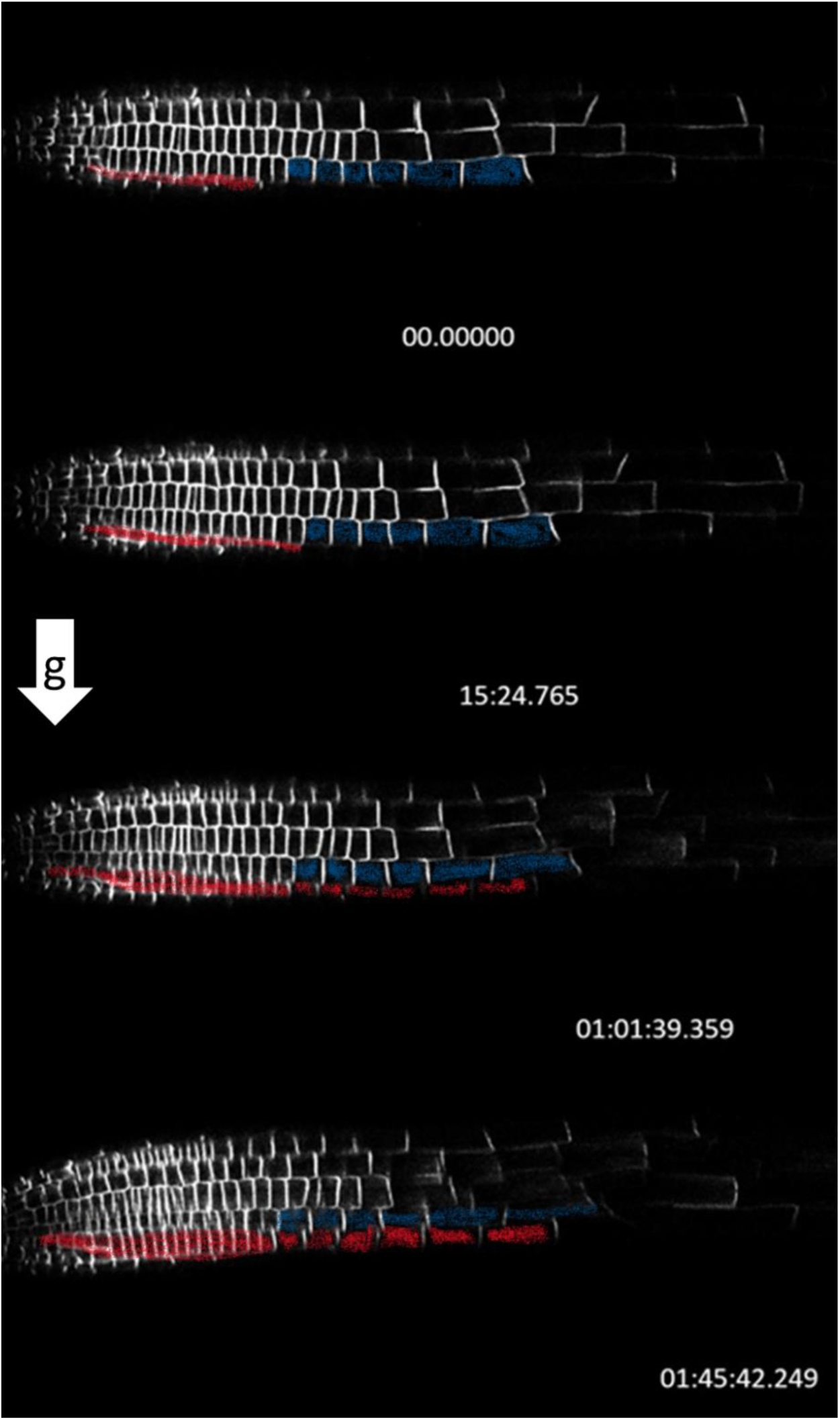
Gravitropic response seems to require epidermal cell file rotation starting at the very root tip towards the elongation zone, where in a neighboring cell file few cells alter their cell expansion to initiate the curvature during directional root growth adaptation.

It will be interesting to investigate if re-orientation of the root tip during the gravitropic response occurs because of the modulation of cellular expansion of the LRC, which may result in a pulling movement and thereby corrects the path of the root tip. Or how the sliding of the epidermal cell file is initiated by mechanoadaptation already in the meristematic zone and which cell mechanoproperties are required to ensure the transmission into a twisting movement towards the elongation zone. The opposing movements of individual cell files force few cells to change their cell expansion/occupancy, which results in the curvature needed for directional growth adaptation. Roots are exposed to everchanging growth conditions, and it is known that manifold abiotic and biotic stimuli are altering the ability of signal transmission between meristem and elongation zone, and also alter mechnaoproperties of epidermal root cells ^[25–30]^.

## Material and Methods

### Plant material and growth conditions

*eir1-4 PIN2::PIN2:mcherry* and *Arabidopsis thaliana ecotype Columbia* expressing PIN2:GFP were used in this study. Seeds were surface sterilized for 5 min using 10% (v/v) bleach containing 0.001% Triton X-100, washed with sterile water and then stratified at 4 °C for 48h in the dark. Seeds were germinated on media containing half-strength Murashige and Skoog (MS) media, supplemented with 1% agar, pH: 5,8. It is important to mention that no carbon source was added to the medium. Seedlings were grown vertically for 6 or 7 days under a continuous temperature of 21 °C with a 16-h photoperiod (100μmolm−2 s−1) using the D-root system ^[16]^.

### Plant imaging

For confocal microscopy the seedlings were transferred to a chambered coverslip according the Sandwich method ^[23]^. Imaging was performed with a Zeiss880 confocal microscope equipped with a vertical stage ^[22]^ with ability to rotate the chambered coverslip to stimulate gravitropic response. Images were taken via objective Zeiss Plan-Apochromat 20x/0.8. Images were acquired by using Zen Black (Zeiss) and processed using Zen Black and Blue.

## Acknowledgements

I want to thank Jozef Lacek for assistance in the lab and the Imaging Facility of the Institute of Experimental Botany AS CR supported by the MEYS CR (LM2018129 Czech-BioImaging) and IEB AS CR. This research was funded by the Ministry of Education, Youth and Sports of Czech Republic from European Regional Development Fund ‘Centre for Experimental Plant Biology’: Project no. CZ.02.1.01/0.0/0.0/16_019/0000738 and the Czech Science Foundation (19-13375Y).

